# Seizure Event Detection Using Intravital Two-Photon Calcium Imaging Data

**DOI:** 10.1101/2023.09.28.558338

**Authors:** Matthew A. Stern, Eric R. Cole, Robert E. Gross, Ken Berglund

**Author notes:** Authors Contributed Equally (Matthew A. Stern, Eric R. Cole). Address all correspondence to Ken Berglund.

## Abstract

**Significance:** Genetic cellular calcium imaging has emerged as a powerful tool to investigate how different types of neurons interact at the microcircuit level to produce seizure activity, with newfound potential to understand epilepsy. Although many methods exist to measure seizure-related activity in traditional electrophysiology, few yet exist for calcium imaging.

**Aim:** To demonstrate an automated algorithmic framework to detect seizure-related events using calcium imaging – including the detection of pre-ictal spike events, propagation of the seizure wavefront, and terminal spreading waves for both population-level activity and that of individual cells.

**Approach:** We developed an algorithm for precise recruitment detection of population and individual cells during seizure-associated events, which broadly leverages averaged population activity and high-magnitude slope features to detect single-cell pre-ictal spike and seizure recruitment. We applied this method to data recorded using awake *in vivo* two-photon calcium imaging during pentylenetetrazol induced seizures in mice.

**Results:** We demonstrate that our detected recruitment times are concordant with visually identified labels provided by an expert reviewer and are sufficiently accurate to model the spatiotemporal progression of seizure-associated traveling waves.

**Conclusions:** Our algorithm enables accurate cell recruitment detection and will serve as a useful tool for researchers investigating seizure dynamics using calcium imaging.

## 1. Introduction

Epileptiform activity in the brain has traditionally been treated and studied using electrophysiological recording methods, such as scalp electroencephalography (EEG) and local field potential (LFP) recordings. However, these recording modalities lack the spatial resolution and cellular specificity needed to understand how different types of neurons interact at the microcircuit level to produce aberrantly synchronous activity.^1^ Calcium imaging is a promising technique that can overcome these limitations in the study of seizure dynamics, providing superior spatial and cellular resolution that could lead to improved understanding and therefore more spatiotemporally precise treatment of seizures.^2^

While a dense body of literature is available describing the characteristics of epileptiform activity in LFP recordings, as well as automated methods to detect these events from electrophysiological data, these are understudied for optical imaging modalities. Intravital calcium imaging data contains largely different sources of noise and signal-to-noise characteristics and thus requires novel solutions for identifying, extracting, and modeling seizure dynamics in order to make the use of this technique accessible, reproducible and standardized in the field of epilepsy research. While event detection approaches exist for calcium imaging, they have largely been limited to traditional domains of systems neuroscience where repeated trials can be leveraged to minimize the sources of noise. However, calcium imaging in epilepsy models poses a unique challenge for automated event detection in that seizure events are highly irregular in their timing, can express with a wide variety of different waveform patterns, and must be detected on single trials, thus necessitating more novel approaches.

In this study, we demonstrate an accurate and computationally efficient signal processing strategy to identify seizure-related events in calcium imaging data – specifically the timing of pre- and inter-ictal spikes, seizure invasion, and propagating waves associated with seizure termination at both the population level and that of individual neurons. We validated this method using two-photon calcium recordings collected from a mouse pentylenetetrazol (PTZ) model of generalized seizures, quantified its performance against manual labels provided by an experienced researcher, and showed that the resulting event times were sufficient to accurately model spatiotemporal dynamics of the seizure associated traveling waves. A MATLAB repository with code for this method and its evaluation was made available online to improve and standardize the study of epilepsy using calcium imaging.

## 2. Methods

### 2.1. Experiments and data collection

#### 2.1.1. Cranial window surgery

All the procedures involving live animals were conducted with approval from and in accordance with Emory University Institutional Animal Care and Use Committee. Stereotactic cranial window surgery and intracortical delivery of adeno-associated virus (AAV) was performed on adult (≥ 90 days old) VGAT-Cre mice (Slc32a1-IRES-Cre, Jackson Laboratory, Stock No. 028862) backcrossed to C57BL6/N background (*N*=4). Electrodes for simultaneous EEG recording and headplates to facilitate awake head fixed recording were placed. The procedure was adapted from standard protocols for concentric cranial window implantation.^3, 4^ In brief, mice were secured in a stereotaxic frame and maintained under anesthesia (1.5% isoflurane balanced in oxygen (1L/min)). A 3 mm craniotomy was performed over the primary motor cortex. Stereotactic injections of AAV (500 nL; 2nL/s) encoding the genetically encoded calcium indicators (GECI) jRGECO1a^5^ or jYCaMP1s^6^ were performed through a pulled glass capillary tube (Nanoject 3.0, Drummond) at 300*μ*m and 600*μ*m deep to the pial surface (0.30 mm anterior and 1.75 mm lateral to Bregma).^7, 8^ AAVs were produced in house following standard procedures.^9^ A thin polyamide insulated tungsten wire electrode with exposed tip (125*μ*m; P1Technologies) was then placed intracortically at the posteromedial edge of the craniotomy, with ipsilateral reference and contralateral ground stainless steel screw electrodes (E363/96/1.6/SPC, P1Technologies) placed in the skull over the cerebellum in 0.7 mm burr holes. All electrodes were prefabricated with gold pins to facilitate easy preamplifier attachment for recording. A concentric window (inner diameter 3 mm, outer diameter 5 mm; D263 #1 thickness coverglass, Warner Instruments) was then placed plugging the craniotomy. The window and electrodes were affixed to the skull using dental acrylic (C&B Metabond, Parkell) along with a stainless steel headplate (Neurotar). The skin was closed to this headplate using tissue adhesive (Vetbond, 3M).

#### 2.1.2. Intravital awake two-photon calcium imaging

Resonant scanning (30 Hz, 512 x 512 pixels) two-photon imaging was performed on mice during acute induced seizures beginning one month following surgery to allow for adequate GECI expression. For each imaging session, mice were head-fixed in a carbon fiber airlifted chamber (Mobile HomeCage, Neurotar) and positioned under a long working distance 16x objective (water-immersion, N.A. 0.80, Nikon). EEG electrodes were connected to an AC preamplifier and a data acquisition system (Pinnacle Technologies). Imaging was performed using a two-photon microscope (HyperScope, Scientifica) equipped with a pulsed tunable infrared laser system (InSight X3, Spectra-Physics) and ScanImage (Vidrio Technologies) controller software. GECI fluorescence was excited using a 1030 nm wavelength light and fluorescence emission was separated by a dichroic mirror (565LP, Chroma) with band pass filters (ET525/50m-2p and ET620/60m-2p, Chroma) and collected using GaAsP and multi-alkali red-shifted PMTs, respectively. All imaging data was synchronously collected with EEG (2 kHz sampling rate). Each recording session began following subcutaneous injection of pentylenetetrazol (40-50 mg/kg; P6500, Millipore-Sigma; sterile saline diluent) and lasted 20-45 min depending on the course of the seizure. Multiple recordings were performed in mice for whom the seizures were not fatal, with sessions separated by at least a week. Data from a representative recording is presented in Fig. 1.

**Fig. 1:**
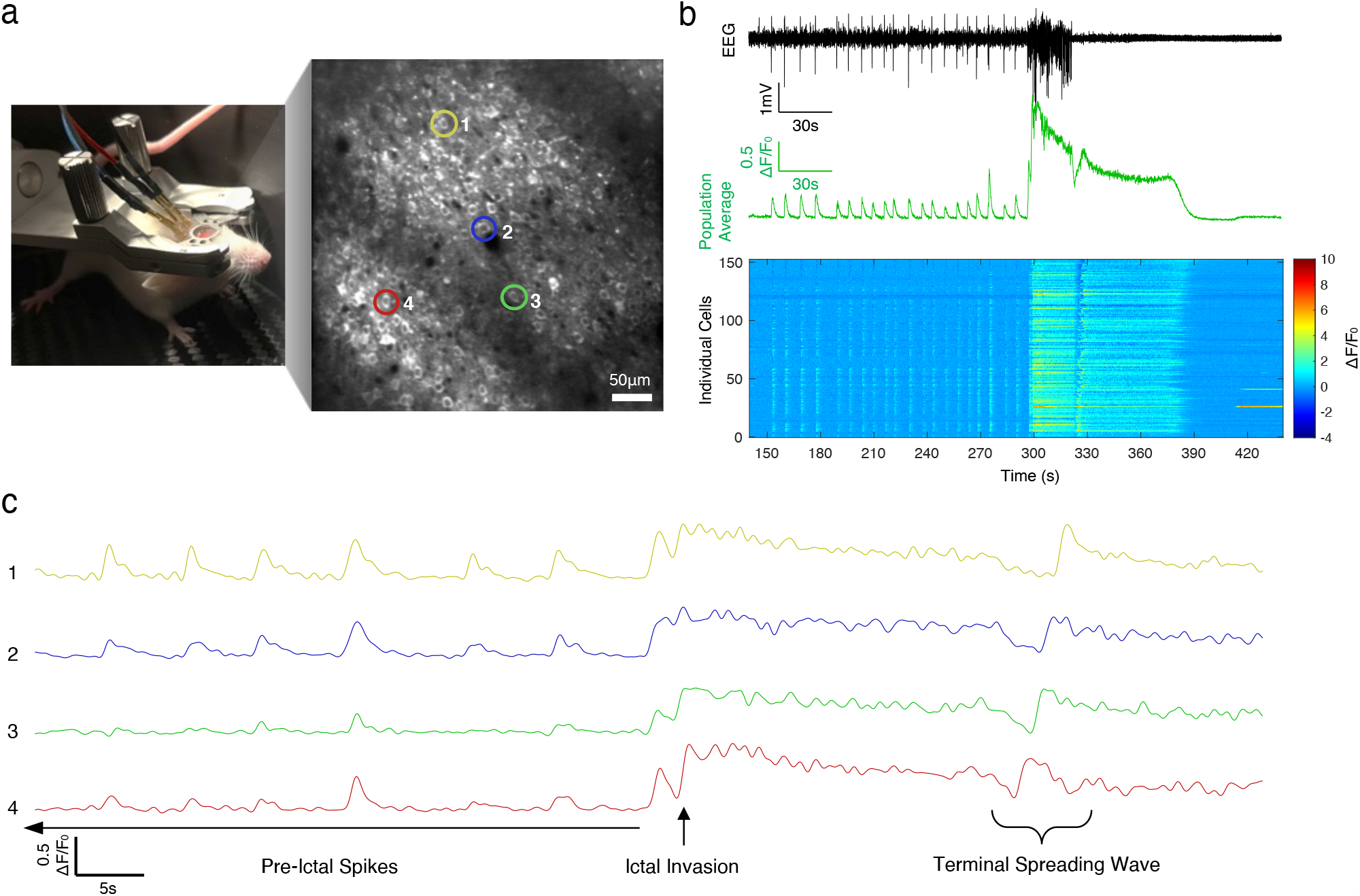
Two-photon calcium imaging during pentylenetetrazol-induced seizures. a) Image of mouse head-fixed in air-suspended chamber for awake intravital two-photon imaging with simultaneous EEG (objective lens removed) and a time-averaged projection of a representative imaging field of view (cortical layer 2/3; depth: 150 *μ*m from pial surface) of cells expressing jYCaMP1s. ROIs highlighted of representative individual cells spanning the field. b) Mean cell calcium fluorescence (green) with synchronized EEG recordings (black) during a PTZ induced seizure with corresponding raster of individual cell calcium transients (y-axis: neurons ordered from left to right across the field). c) Normalized calcium fluorescence traces(ΔF/F_0_) of representative cells indicated in panel a during the three phases of seizure activity from which event times will be extracted (pre-ictal spikes, ictal-invasion, terminal spreading wave).

### 2.2. Description of event detection algorithm

The following section, as outlined in Fig. 2, describes the approach our algorithm uses to determine the population event times and individual cell recruitment to the events, such as preictal spikes and calcium waves during and after a seizure, using two-photon calcium imaging data.

**Fig. 2:**
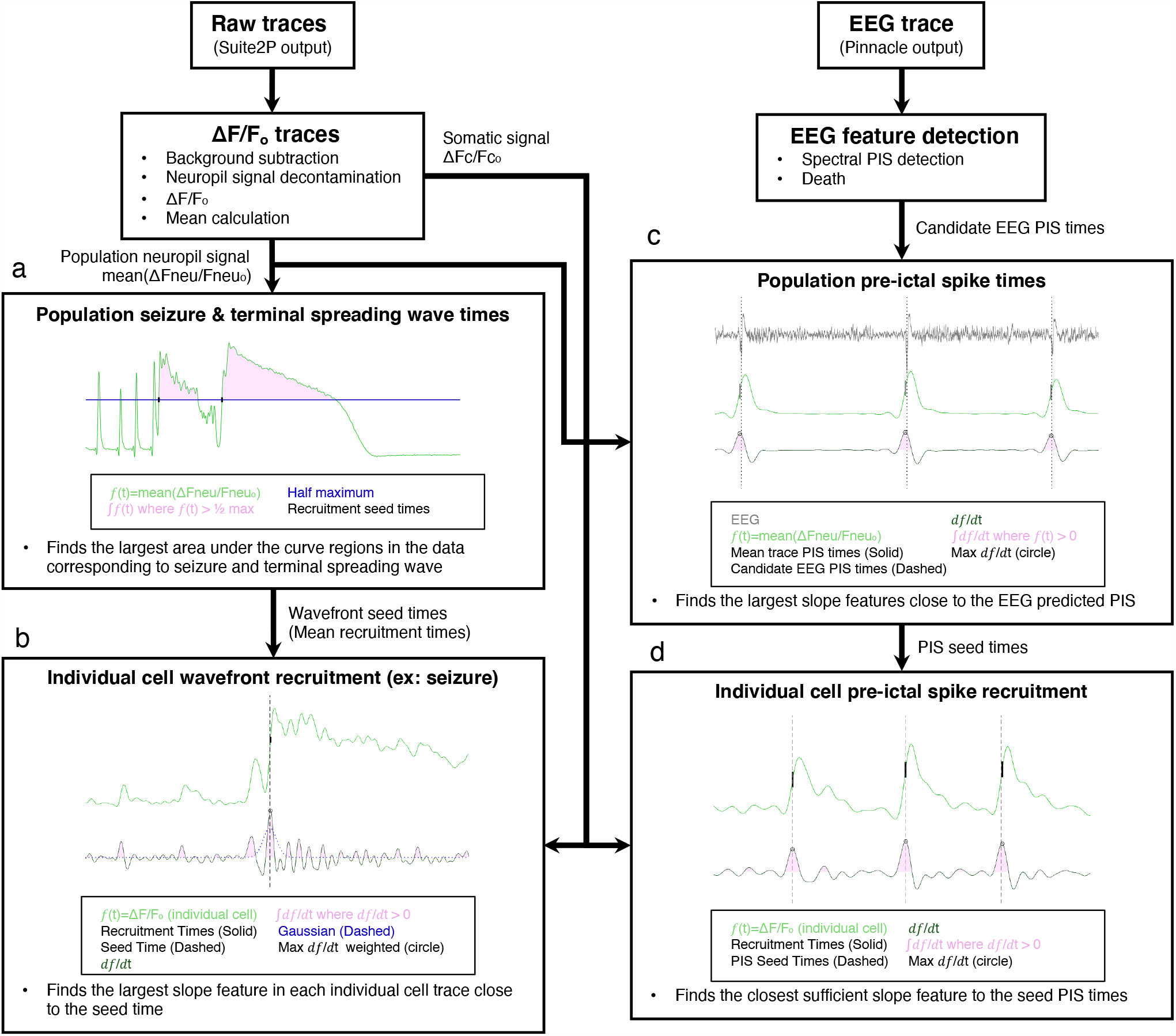
Algorithmic flowchart for detecting seizure-related events in calcium imaging data. This flowchart demonstrates how raw calcium transient data and EEG are processed to determine preictal spike and wavefront event times. The calcium transient output from suite2p is first used to generate mean event times using different methods tailored to the event type (pre-ictal spikes (PIS), seizure or terminal spreading wave). These mean event times are then used as seeds to guide event detection for individual cells. a) i) Integrate *f*(t) where *f*(t) > *½* max. ii) Select two largest events as seizure and terminal spreading wave; use the first *½* max intercept time point as each event’s recruitment time. b) i) Integrate *df*/*d*t for all contiguous regions where *df*/*d*t > 0. ii) Multiply each slope-integral feature by its Gaussian-weighted time difference from the mean seizure seed time; then for the largest event, select the time point of max *df*/*d*t. iii) Define each cell as recruited if its recruitment is within 3s of the mean, and there is a > 20% post-recruitment f(t) increase. c) i) Compute slope-integral features (as in b(i)). ii) Select top 15% of events with highest feature value for further analysis. iii) Discard events with a low threshold feature value or a recruitment time (point of max *df*/*d*t) >0.2s from the closest EEG-detected spike. iv) Generate EEG SWD kernel to find missing events. d) i) Compute slope-integral features. ii) Select top 20% of events with highest feature value for further analysis. iii) Select the event closest to the population PIS seed time. iv) Discard low peak-height events (<4 standard deviations from the mean) that are > 1.5 seconds away from PIS seed time.

#### 2.2.1 Data pre-processing

Motion registration, region of interest (ROI) detection and calcium transient extraction of soma and surrounding neuropil were performed using the Suite2P software package^10^ with integrated Cellpose.^11^ The raw fluorescence traces were background subtracted taking the global minima of the neuropil transients for each recording as a proxy for the background signal. A clean somatic signal was then generated by subtracting 70% of the surrounding neuropil signal from the somatic signal to remove out-of-plane contamination.^10^ This step also serves to adjust for any photobleaching that occurs in the indicators. While this is not typically a problem for cytosolic indicators, any fluorophore bound or tethered to the membrane can suffer some photobleaching as it remains in the focal plane over long imaging sessions. These traces were then normalized as ΔF/F_0_, using the mean of the first 30 seconds of recording as baseline fluorescence (F_0_).

#### 2.2.2. Pre-ictal spike recruitment detection

To identify the times of individual cell recruitment to pre-ictal spikes, global pre-ictal spike times were determined using the averaged calcium trace of each cell’s nearby neuropil signal and the EEG recording. These were then used as “seed times” to guide the identification of individual cell firing times during each mean spike event. We found that preictal spikes were more prominent in both space and time in the neuropil signal, therefore providing better estimation of mean activity than with individual cells’ activity in the soma to guide individual cell event detection.

To determine the global pre-ictal spike times, we began by detecting spike wave discharges on an EEG recording and verified their agreement with spike times determined from the large population mean calcium spikes. For the EEG spike detection, a spectrogram was computed using Welsh’s power spectral density estimate periods of high power near theta (3-15 Hz) relative to power near low gamma (20-55 Hz) power that were then followed by a period of low gamma. EEG spikes were indexed by the peak of the spike on the spike wave discharge, being the most prominent and least variable feature of the spike. To detect calcium spikes, the mean calcium trace was first double-reverse filtered using an adaptive filtering routine (1 Hz, Butterworth lowpass filter, order 3 to 5) that preserves the phase of the signal. All segments of the trace with positive slope were isolated and then a magnitude feature for each segment was computed as the integral of the first derivative of the signal (the “slope-integral” feature) to account for steepness of slope, peak height, and rise-time duration. The periods of the mean calcium trace that corresponded to the top 15% of these were then thresholded by taking the integral of the periods and comparing it to the integral of a model spike (gaussian with a peak height of the standard deviation of the trace and a standard deviation (σ) of 0.33 s to give a 1s width at 1.5 σ) representative of a typical spike weight under normal physiologic conditions. The exact time of each calcium spike was determined by its point of steepest slope. These spikes were then cross-referenced with those detected on EEG, so that only a pair of spikes occurring within 0.2 seconds on each modality was included in the final set of pre-ictal spike times. Then, the time observed on the EEG was used as the recruitment time for each spike.

While false positives are typically virtually eliminated after applying the above steps, we sought to further minimize false negatives for spikes of low amplitude but that were nevertheless physiological. To do this, a spike wave discharge kernel was generated by taking the mean of the EEG for each calcium and EEG concordant pre-ictal spike determined thus far in the set. This kernel was then used to find the points of greatest convolution in the EEG trace, finding previously missed pre-ictal spike based on their waveform similarity to the physiological spikes. This step ensures to include isolated spikes that may not be necessarily reflected as population spikes in calcium.

In order to next determine the points of individual cell recruitment to pre-ictal spikes, all individual clean somatic traces were low-pass filtered to 1 Hz using the same method as the mean trace. As was done in the mean trace, all regions of each individual trace with positive slope were isolated and then the magnitude of that slope was computed as the integral of the first derivative of the signal (the slope-integral feature). These events were then indexed by the point of steepest slope. The top 20% of these events were taken and the events closest to each of the pre-ictal spike seed times was determined. A cell was considered recruited if this time fell within 1.5 s of the seed time and the peak of the calcium event was greater than 4 σ above the cell’s mean.

#### 2.2.3 Wavefront recruitment detection

Detection of seizure wavefront and terminal spreading wave at the population level was performed using the mean neuropil calcium trace, which provided better detection than using the mean somatic calcium trace, likely because the neuropil signal was less variable across cells than the somatic signal. This signal was first low pass filtered to 1 Hz using the same filtering routine as used in the pre-ictal spike detection. The integral of the trace was then taken of all contiguous intervals above a half maximum and the two largest events were determined. The first of these was taken as the seizure event and the second of these was taken as the terminal spreading wave event. These were indexed by the first time point where the trace crossed the half maximum.

The wavefront event time was then used as a seed to guide individual cell recruitment detection. As in pre-ictal spike detection, all individual clean somatic traces were low pass filtered to 1 Hz, and all regions with positive slope were segmented, with the magnitude of the slope computed as the integral of the first derivative of the signal (slope-integral feature). These events were then indexed by the point of steepest slope. The slope magnitudes were then weighted by a Gaussian distribution centered at the mean event seed time, in order to avoid false-positive detection of events more temporally distant from the seed time. To ensure detection of the seizure wavefront and not the sentinel spike often occurring immediately ahead of the wavefront, a narrow gaussian (σ=1 s) was used, while a larger Gaussian (σ=5 s) was used for the terminal spreading wave given its much slower propagation. The period with the maximum weighted slope-integral feature was then taken as the individual cell’s recruitment time. A cell was considered recruited if there was at least a 20% increase in average signal for that cell before and after recruitment and the time detected was within 3 s of the seed time for seizure wavefront and within 5 s for the terminal spreading wave. The before and after periods to account for signal change were taken as 10 s for seizures given the occurrence of pre-ictal spikes and to leverage the large and sustained signal change that occurs in the cells during seizure. For the terminal spreading waves the best detection period was a narrower window of 5 s due to the higher signal variability between the seizure and terminal wave. The terminal spreading waves invaded the imaging field of view at different time points between seizures, with some entering during the final phase of ictal activity and others entering after ictal activity had largely died out, contributing to variable calcium levels at the begging of the events.

Note that in some instances (*N*=2), the seizure was fatal and under these circumstances the recording was truncated shortly after the point of death for event detection purposes.

### 2.3 Algorithm Evaluation

To evaluate the algorithm’s performance, an expert blind to the automated detection algorithm described above used a MATLAB-based GUI (Fig. S1) to visually inspect the ΔF/F_0_ traces and manually mark the times of pre-ictal spikes, seizure invasion and terminal spreading wave events of activity averaged across the population and of individual neurons (detected events shown in Fig. 3a for pre-ictal spikes and Fig. 4a for seizure events). The times for all event types (pre-ictal spikes, ictal and post-ictal waves) was defined as the point of maximum slope. While labeling, the expert was able to view the EEG recording and averaged population ΔF/F_0_ as an additional reference to help decide whether candidate events were seizure-related instead of unrelated noisy fluctuations in the signal.

**Fig. 3:**
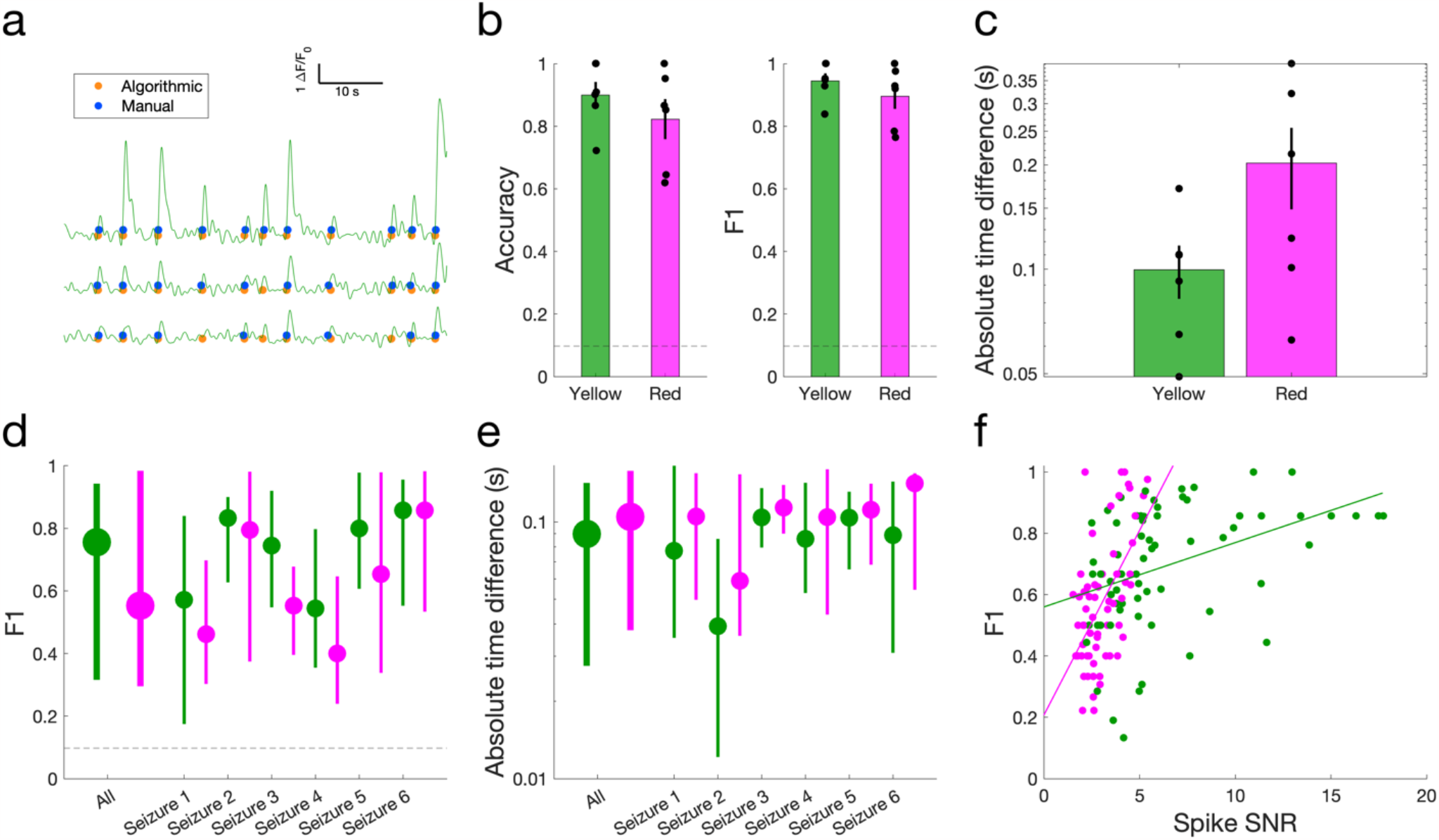
Pre-ictal spike recruitment detection performance. a) Example manually (blue) and algorithmically (orange) detected pre-ictal spike events for three cells within 60 seconds before seizure invasion. b) Accuracy and F1 score quantifying algorithmic performance in detecting preictal spikes in averaged *Δ*F/F_0_ signal from both yellow (green) and red (magenta) cell populations across 6 seizures. Dashed line: random-chance performance (0.098). c) Average absolute time difference between manually and algorithmically detected pre-ictal spikes on averaged *Δ*F/F_0_. d) F1 score quantifying algorithmic performance in detecting pre-ictal spikes of individual cells, shown for all and individual seizures. Circles and bars indicate median and bootstrapped 95% confidence interval. e) Absolute time difference between manually and algorithmically detected pre-ictal spikes of individual cells, shown for all and individual seizures. f) Detection performance vs. cell signal-to-noise ratio of pre-ictal spike amplitude. (Yellow: p = 7.4e-4, R^2^ = 0.162; Red: p = 1.25e-6, R^2^ = 0.318).

**Fig. 4:**
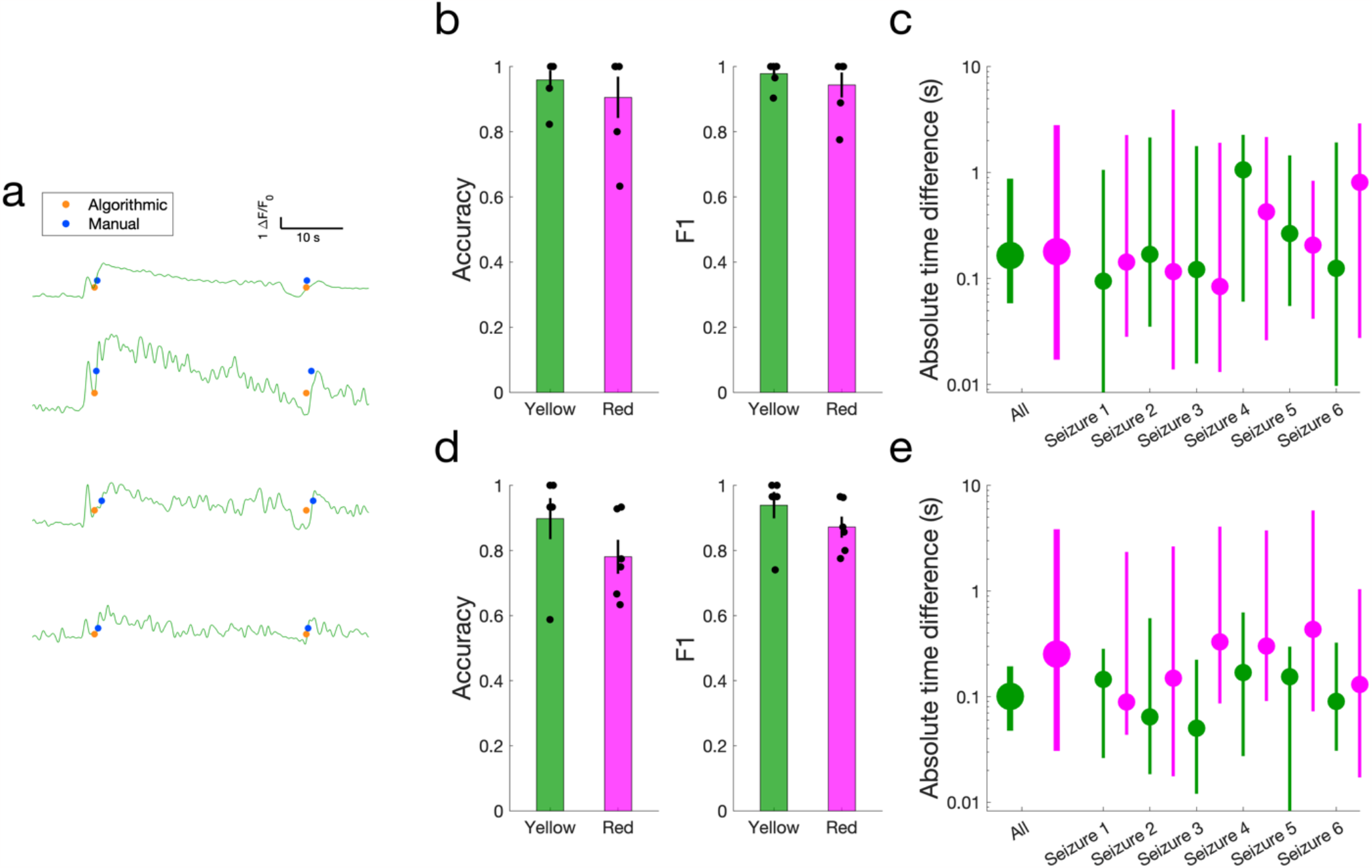
Wavefront recruitment detection performance. a) Example manually detected (blue) and algorithmically detected (orange) seizure invasion and terminal spreading wave events, shown using *Δ*F/F_0_ averaged across the cell population (top) and for three example cells (second through bottom). b) Accuracy and F1 score quantifying algorithmic performance in detecting seizure invasion for individual cells from both green (green) and red (magenta) cell populations. c) Average absolute time difference between manually and algorithmically detected seizure invasion events for individual cells, shown for all and individual seizures. d) Accuracy and F1 score quantifying algorithmic performance in detecting seizure terminal spreading wave events for individual cells from both green (green) and red (magenta) cell populations. e) Absolute time difference between manually and algorithmically detected seizure terminal spreading wave events for individual cells, shown for all and individual seizures.

We computed accuracy metrics between detected and manual labels by iterating through manually detected events and searching for a matching auto-detected event within 1 second, producing a true positive detection. Any unmatched manual event was designated as a false negative and any unmatched algorithmic event as a false positive. We computed the accuracy and F1 score (geometric mean of precision and recall – *i.e*., positive predictive value and sensitivity) for each event type:

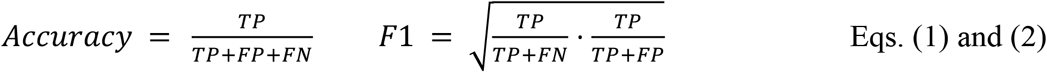

Where TP is the true positive count, FP is the false positive count, and FN is the false negative count.

When interpreting these metrics, it should be noted that the true negative rate is ill-defined due to the sparsity of seizure events; *i.e*., the algorithm can easily properly detect that there is no event during most of the duration of the recording, but this result is trivial and properly quantifying it is an ambiguous problem. Therefore, the results should be interpreted in two ways. First, we consider the F1 score to be a better evaluation metric than accuracy because its computation is naturally independent of the true negative rate. Second, 50% accuracy is no longer an appropriate baseline corresponding to random chance-level performance; rather, true random-chance event detection performs much worse than 50%.

To provide a statistically accurate performance baseline, we simulated a Poisson process and quantified the resultant F1 score when randomized event times are used for detection. For each recorded seizure, we computed the average frequency of spikes in the pre-ictal period, then used the reciprocal of this value as the average Poisson distribution inter-event time interval to generate randomized event times for the duration of the pre-ictal period. We then computed accuracy and F1 metrics between these randomized event times and manually identified times in the same manner as above. We repeated this simulation 50 times per seizure and report an overall mean F1 score of 0.098. Pre-ictal signal-to-noise ratio (SNR) was calculated for each cell by computing the mean height of all peaks with greater than 1 ΔF/F_0_ prominence during the pre-ictal period, then dividing by the variability of baseline activity during this period (quantified by 40^th^ percentile minus 10^th^ percentile of the time series, so that the noise estimate excludes outliers and data values of the peaks themselves). Linear regression was used to demonstrate the relationship between the SNR of the data and performance of our algorithm.

## 3. Results

We performed simultaneous *in vivo* two-photon calcium imaging and EEG recording in an awake, head-fixed mouse with prior injection of AAV for expression of a GECI in the motor cortex (Fig. 1a). A seizure was elicited by an injection of PTZ (40-50 mg/kg, i.p.), resulting in sporadic preictal spikes for up to 10 minutes followed by a seizure manifested as a rapid invasion of calcium wave into the field of view. Often times, shortly after the cessation of the seizure, we encountered a much slower propagation of calcium wave (Fig. 1b). We developed an algorithm to automatically detect these different event types (Fig. 2). This algorithm can detect calcium transients even in noisy recordings by utilizing population responses (EEG and population-averaged calcium transients) as seed-times to guide detection of high-slope events in individual cells.

### 3.1. Pre-ictal spike detection

We first evaluated our method for detecting pre-ictal spikes by comparing its performance against manual labels, measuring the accuracy and absolute time-difference between algorithmic and manual event detection (*N*=6 recordings). When applied to the population mean ΔF/F_0_, our method achieved 0.900±0.042 (mean ± SEM; *n*=6) accuracy and 0.945±0.024 (mean ± SEM; *n*=6) F1-score in detecting preictal spikes using the yellow indicator (Fig. 3b). Using the red indicator, accuracy was 0.823±0.064 (mean ± SEM; *n*=6) and F1 was 0.896±0.040 (mean ± SEM; *n*=6). The average time difference between automatically detected events and manually labeled events (Fig. 3c) was 0.0996±0.018 s (mean ± SEM; *n*=6; yellow) and 0.202±0.0536 s (mean ± SEM; *n*=6; red), considerably less than the typical pre-ictal spike width and comparable to 3-7 imaging frames (frame rate: 30 Hz). In the detection of individual cell events, the algorithm demonstrated an overall median F1-score of 0.755 (yellow; 95% confidence interval = [0.321, 0.943]; *n* = 72 cells) and 0.553 (red; 95% confidence interval = [0.293, 0.983]; *n* = 72 cells) in detecting spike events, with a median time difference of 0.090 (yellow; 95% CI = [0.027, 0.142]; *n*=72) and 0.105 (red; 95% CI = [0.038, 0.158]; *n*=72;) . Worst-case performance for any individual cell from any seizure (via lower bound of the 95% confidence interval) was consistently greater than the random-chance baseline of 0.095 estimated via a simulated Poisson process. Overall, accuracy was higher with the yellow indicator because of its generally higher signal-to-noise characteristics. F1 score for preictal spike detection increased as the cell’s spike signal-to-noise ratio improved (Fig. 3f).

### 3.2. Seizure invasion and activity at termination detection

We next quantified and compared the performance of our algorithm in detecting seizure invasion and activity at seizure termination (Fig. 4). For detecting seizure invasion events using the yellow indicator, the algorithm achieved a detection accuracy of 0.960±0.029 (mean ± SEM; *n*=6) and F1 score of 0.978±0.016 (mean ± SEM; *n*=6) in predicting which cells were recruited into the seizure, with a median absolute time difference of 0.165s (95% CI = [0.058, 0.882]; *n*=96 cells). Use of the red indicator yielded accuracy of 0.906±0.064 (mean ± SEM; *n*=6), F1 score of 0.944±0.0383 (mean ± SEM; *n*=6), and a median time difference of 0.179 s (95% CI = [0.017, 2.808]; *n*=96 cells).

For detecting terminal seizure events using the yellow indicator, we achieved a detection accuracy of 0.898±0.063 (mean ± SEM; *n*=6), and F1 score of 0.940±0.052 (mean ± SEM; *n*=6), in predicting which cells were recruited into the seizure, with a median absolute time difference of 0.101 s (95% CI = [0.047, 0.194]; *n*=96 cells). Use of the red indicator yielded accuracy of 0.781±0.052 (mean ± SEM; *n*=6), F1 score of 0.872±0.033 (mean ± SEM; *n*=6), and a median time difference of 0.253 s (95% CI = [0.031, 4.012]; *n*=96 cells).

As was the case in the pre-ictal detection, the performance of the algorithm was somewhat contingent on the indicator being used, the relative SNR of the indicator, and the dynamics of the cell populations being recorded. Nevertheless, our algorithm achieved detected recruitment times concordant with those of an expert scorer across indicators for both the seizure wavefront and terminal spreading wave.

### 3.3. Seizure wavefront modeling

To illustrate the utility of this detection method, we calculated the rates and directions at which calcium waves propagate during and after a seizure using the position of the cells and the automatically detected recruitment times. We applied a spatial linear regression algorithm (originally developed to model interictal events in human intracranial data)^12, 13^ to our detected recruitment times to model propagation of the seizure wavefront throughout the imaged cortex in one example recording. Fig. 5a shows sequential imaging frames during a seizure invasion event (progressing from bottom-right to top-left; top) and a terminal spreading wave (progressing from left to right; bottom). Recruitment time was automatically determined for each cell and visualized in Fig. 5b. In combining these techniques, we were able to estimate statistically significant propagation vectors (seizure: *p* <0.001, velocity = 421*μ*m/s, angle = 0°; terminal spreading wave: *p* <0.001, velocity = 68*μ*m/s, angle =152°) for each traveling wave that appears to be consistent with direction and velocity with the sequential frames. Therefore, our automated detection method was demonstrated to be sufficiently accurate to study the roles of different cell types and individual neurons in seizure activity.

**Fig. 5:**
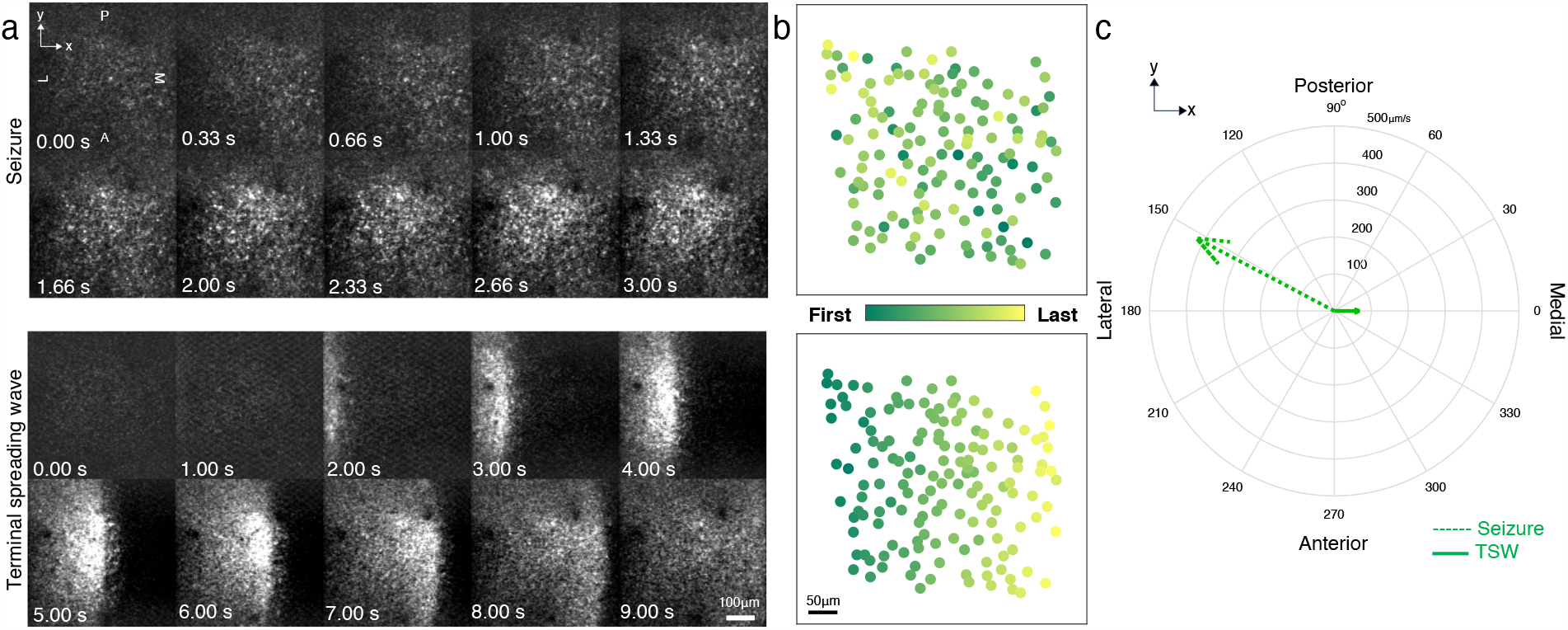
Modeling spatiotemporal dynamics of wavefront propagation. a) Calcium imaging frames demonstrating wave propagation during ictal-invasion (top) and terminal spreading wave (bottom) events. b) Heatmap of determined recruitment times for identified cells during the events shown in panel a, with color corresponding to recruitment order. c) Propagation vectors modeled by applying spatial linear regression to the neuronal recruitment times shown in panel b, showing wavefront direction (vector angle) and propagation speed (vector magnitude).

## 4. Discussion

In this study, we demonstrated an accurate and computationally efficient signal processing strategy to identify pre-ictal, ictal, and post-ictal events using calcium imaging data. We showed that this method can achieve high accuracy for each of these event types and that the precision of the identified event times is sufficient to map the spatiotemporal progression of various seizure-related traveling waves, including seizure wavefronts and various terminal spreading waves, including purportedly cortical spreading depression and hypoxia induced waves of death. In the study of epileptic microcircuit dynamics, the ability to determine exact recruitment times for individual cells is of great importance, particularly to evaluate the changes in population synchrony and patterns of propagation. This in turn enables the study of questions at the forefront of the field, such as the role of inhibition in either restraining the network (inhibitory restraint) or paradoxically synchronizing it (inhibitory rebound).^14, 15^

Our method has been informed by several approaches utilized by other groups. Principally, for defining the time point of recruitment, we use the local timepoint of maximum slope to determine when the cell or population is recruited into epileptiform activity.^16-18^ Other groups have used the time at which the signal exceeds a threshold value such as multiple standard deviations above baseline activity, crossing the half-maximum point of a trace or exceeding a percentage increase from baseline.^19-27^ One group made this method robust by using a moving window filter to detect high-threshold events, and used the duration where the event was above threshold to differentiate seizures from spike events.^28^ Still others define pre-ictal and ictal events based solely on the EEG signal and then determine if cells are statistically significantly activated during that event relative to other activity outside the event.^29, 30^ Niemeyer et al. used a clever method for detection of events by first finding global seizure events in the calcium data using a threshold detection in the mean trace and then generated a seed from a global signal and anchoring it to a point of recruitment, being the elbow of trace, or rather the maximum second derivative.^31^

We believe our approach offers several advantages over methods employed by other groups. While another study used lowess smoothing to remove noise before detection^16^, our use of a zero-phase low-pass filter removed shot noise equally effectively without distortion of the signal in time and with orders-of-magnitude lower computational demands. Our method also allows for automatic detection of pre-ictal, ictal, and post-ictal events without needing to manually define which periods of the signal used for which type of analysis. For multiple different event types, we used a slope-integral feature to detect events; we found this feature to be more reliable at isolating events compared to threshold-based or simple slope-based detection, as even after applying the temporal filter, many high-magnitude noise fluctuations still provided false positives. The slope-integral feature solved this problem by leveraging the property that real spikes and wavefront events typically had longer rise times than this type of noise, to enable robust detection.

There were several sources of noise in calcium imaging data that guided the design of our approach and introduce challenges for detection. One major limiting factor is the presence of sentinel spikes, a single large amplitude spike that directly precedes the seizure.^32^ These spikes exhibit large population wide calcium activity often within a few seconds of ictal invasion and thus can complicate the identification of epileptic wavefront recruitment. Additionally, signal changes associated with seizure activity, such as sudden movements of the animal body at a seizure onset, can lead to lateral shifts, axial drifts, and fluctuating shifts in the baseline signal (F_0_) that introduce further challenges. While these can be slightly mitigated mechanically with excellent headplate installation and a high rigidity head-fix apparatus, with online z-shift correction or with post hoc motion registration,^10^ convulsive seizures present a uniquely intense circumstance that can sometime exceed the corrective capacity of these measures to completely prevent these artifacts from impacting the recording quality. Furthermore, microscope scanning shot noise, particularly from high frequency resonant scanning, and reduced sampling rates can affect the accuracy of our method. Another limitation is SNR of the indicator used. Although our detection method performed well using the indicators of our choice, we expect an improvement of the accuracy of our approach by utilizing indicators with higher SNR, such as improved versions of GCaMP.^33^ In fact, F1 score improved as SNR of pre-ictal calcium transients in individual cells increased (Fig. 3f). It is also important to note that manual labels, which we used as a ground truth for evaluation, are not perfect. In particular, some events, especially in low-SNR cells, can be visibly indistinguishable from noise fluctuations, leading to variability in visual inspection. Discarding low-SNR cells might be necessary to ensure accurate event detection, although this decision requires careful consideration when the aim of the study is to accurately capture population variability – *i.e*., cell SNR could be related to a physiological variable of interest (e.g. GABAergic subtype) and therefore this consideration could influence results and conclusions.

Genetic calcium imaging is becoming a more widely used technique to study epileptic networks as it affords great advantage over classically used electrophysiological techniques, principally enabling the recording of hundreds of neurons repeatedly with subtype specificity.^1, 2^ However there remain substantive barriers to the more widespread adoption of these methods including technical expertise and computationally efficient and implementable methods. Currently it seems nearly every group analyzing this data uses a different approach to quantify these results and few offer a quantitative analysis or validation of the accuracy of their methods. There is a gap in the tools readily available that are optimized for the unique situations presented by the high frequency synchronous firing occurring during seizure events. Here we offer an approach and associated code to aid not only in closing this gap, but in facilitating the standardization of the analysis across the epilepsy field. However, it is important to note that as there is a lack of standardization amongst the experimental paradigms used, including model system, indicators used and imaging parameters; some modifications may be necessary to tailor this approach to each lab’s unique circumstances, as is often the case with pre-clinical EEG/LFP seizure and spike detection.

In addition to the study of seizure dynamics, accurate methods to detect and model epileptiform activity in calcium imaging could benefit the long-term development of future epilepsy treatments in translational settings. Despite great advancements for management of epilepsy over the past century, nearly one-third of patients’ seizures are still poorly controlled with medication^34, 35^ and surgical treatment options^36-39^ are only effective for a subset of these patients. Pre-clinical optical imaging with cellular resolution might be used to evaluate novel brain stimulation targets for epilepsy with the potential for superior cell-specific modulation,^40-43^ or to provide unbiased evaluation of different stimulation patterns and parameters^44, 45^ for seizure control at scale.

Efficient seizure detection methods might also be combined with mean-field imaging to provide feedback for real-time optimization and control paradigms, as brain stimulation for seizure control may benefit by responding to the dynamics of different cell populations in real-time.^46-48^ In this way, our work may provide a small but important contribution to the development of novel seizure control strategies in animal models, which could inform treatment approaches for patients with medically intractable epilepsy.

## Code, Data, and Materials Availability

The code needed to perform this analysis is available in our GitHub repository at https://github.com/Stern-MA/2PCISz_RecDetect and the data used to evaluate this approach will be made available upon request to the authors.

## Acknowledgments/Funding Sources

This work was supported by NIH F31NS115479 (MAS), R21NS112948 (REG), S10OD021773 (KB), and the Mirowski Family Foundation (REG).

## Biographies

**Matthew Stern** is currently an MD/PhD candidate at Emory University completing his neuroscience PhD under the advisership of Robert Gross. He received his BS degree in biology from Haverford College in 2011. He is interested in the use of functional optical imaging modalities to advance our understanding of the basic mechanisms of neurological disease, in particular epilepsy, and their utility to inform neurosurgical treatments.

**Eric Cole** is currently a PhD candidate in the joint biomedical engineering program at Emory University and Georgia Institute of Technology under the advisership of Robert Gross. He received his BS degree in bioengineering and electrical engineering from Cornell University in 2018. He is interested in developing real-time algorithmic methods for precision brain stimulation.

**Robert Gross** is the MBNA/Bowman Endowed Chair in Neurosurgery at Emory University with appointments in Neurology and the Coulter Department of Biomedical Engineering of Emory/Georgia Institute of Technology. He directs the Emory Neuromodulation Technology Innovation Center and the Translational Neuro-engineering Research Lab. He received his ScB in Neural Science from Brown University in 1981 and his MD and PhD in Molecular Pharmacology from Albert Einstein College of Medicine in 1990 under the advisership of Charles Rubin.

**Ken Berglund** is Assistant Professor of Neurosurgery at Emory University and leads various imaging projects in the Robert Gross laboratory. He completed his BA in 1997 and PhD in 2002 both in Psychology at the University of Tokyo under the advisership of Masao Tachibana and postdoctoral training at Duke University under the advisership of George J. Augustine.

