## Supplemental Figures for "Seizure Event Detection Using Intravital Two-Photon Calcium Imaging Data"

[a]Authors Contributed Equally

### Supplemental Figures

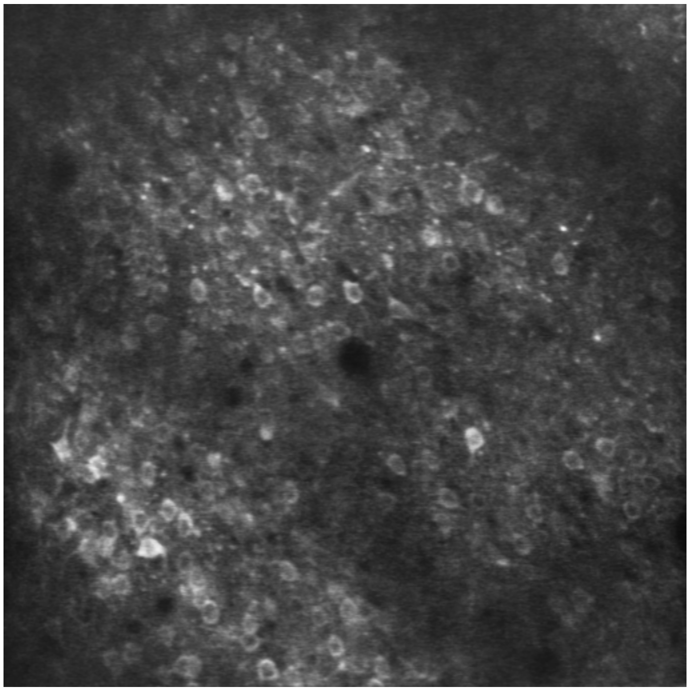

**Fig. S1:** *In vivo* awake two-photon calcium activity (jYCaMP1s) during the progression of a pentylenetetrazol induced generalized seizure from the pre-ictal through the post-ictal phase. Field of view is 450 μm x 450 μm. Playback speed at 4x. (MPEG-4, 20.4MB).

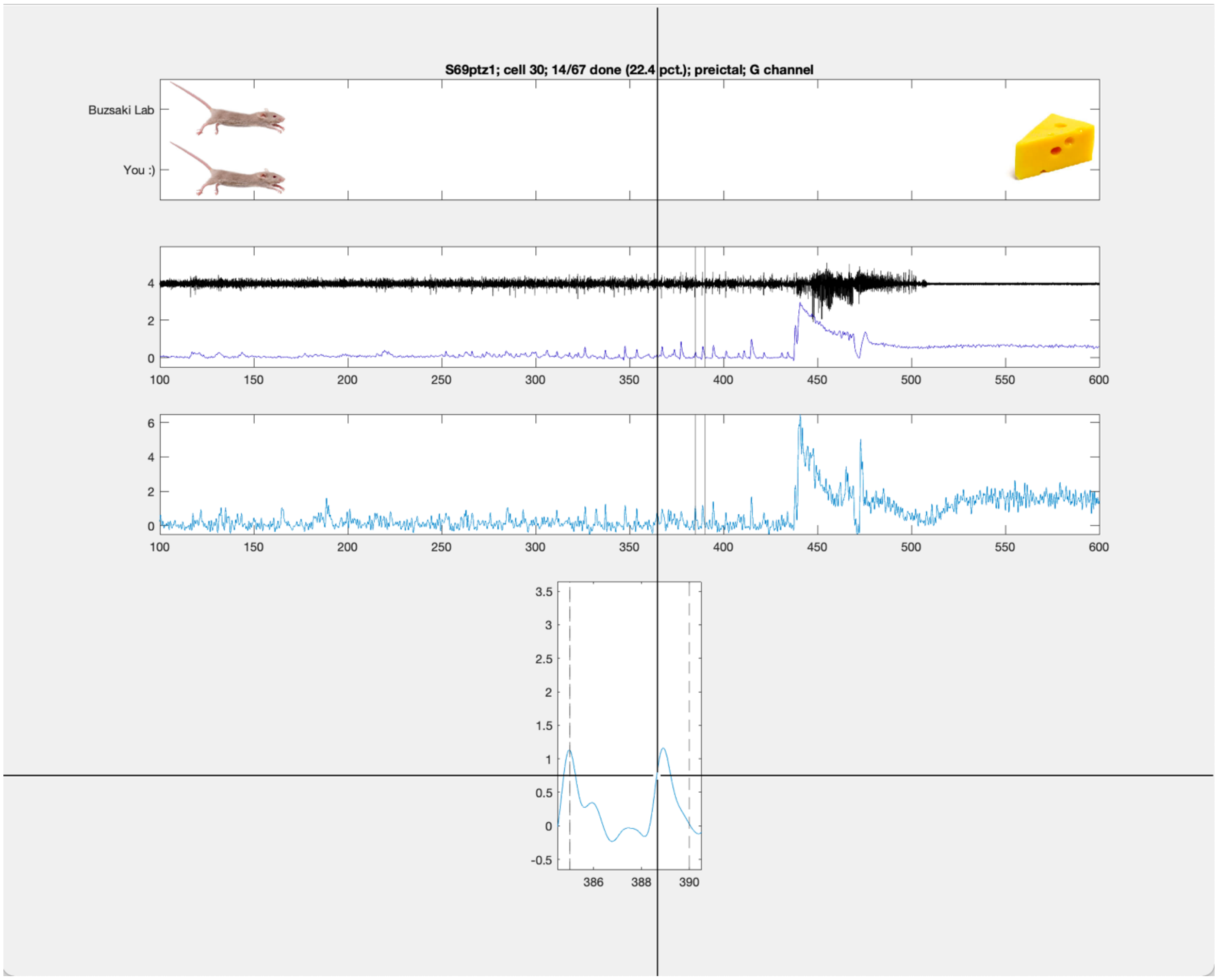


**Fig. S2:** Screen capture of GUI for manual annotation of seizure recruitment events. from top to bottom 1) progress bar 2) reference mean calcium trace and EEG with window bounds indicated depicting region being annotated 3) reference of full individual cell calcium trace 4) annotation window for selection of recruitment points on individual cell trace during a pre-ictal spike with cursor tracking crosshair to use for selection.
